# Tomato yellow leaf curl virus V2 protein plays a critical role in the nuclear export of V1 protein and viral systemic infection

**DOI:** 10.1101/669754

**Authors:** Wenhao Zhao, Yinghua Ji, Shuhua Wu, Elizabeth Barton, Yongjian Fan, Xiaofeng Wang, Yijun Zhou

## Abstract

Geminiviruses are an important group of circular, single-stranded DNA viruses that cause devastating diseases in crops. Geminiviruses replicate their genomic DNA in the nucleus. The newly-synthesized viral DNA is subsequently transported to the cytoplasm, moved to adjacent cells through plasmodesmata with the help of viral movement proteins, and, ultimately, moved long-distance to establish systemic infection. Thus, the nucleocytoplasmic transportation is crucial for a successful infection by geminiviruses. For *Tomato yellow leaf curl virus* (TYLCV), the V1 protein is known to bind and shuttle viral genomic DNA, but the role of V2 protein in this process is still unclear. Here, we report that the nucleus-localized V1 protein dramatically decreases when co-expressed with V2 protein, and that V2-facilitated nuclear export of V1 protein depends on host exportin-α and a specific V1-V2 interaction. Chemical inhibition of exportin-α or a substitutions at cysteine 85 of V2 protein, which abolishes the V1-V2 interaction, blocks the promoted redistribution of V1 protein to the perinuclear region and the cytoplasm. When the V2^C85S^ mutation is incorporated into a TYLCV infectious clone, the TYLCV-C85S causes delayed onset of very mild symptoms compared to wild-type TYLCV, indicating that the V1-V2 interaction and, thus, V2-mediated nuclear export of V1 protein is crucial for viral spread and systemic infection. Our data point to a critical role of the V2 protein in promoting the nuclear export of the V1 protein, likely by promoting V1-mediated nucleocytoplasmic transportation of TYLCV genomic DNA, and in turn, promoting viral systemic infection.

**Author summary:** As both replication and the transcription of geminiviruses occur in the nucleus, transportation of the viral genomic DNA into and out of the nucleus of the infected cells is essential for a successful infection cycle. However, the nuclear export of geminiviruses is still little known and even less is known about the process for monopartite geminiviruses. We use TYLCV, a typical monopartite begomovirus in the family *Geminiviridae*, to examine the nucleocytoplasmic transportation. In this study, we found TYLCV V2 is able to redistribute the nucleus-localized V1 protein to the perinuclear region. Moreover, the nuclear export of V1 protein is dependent on the V1-V2 interaction and host exportin-α. Blocking the V1-V2 interaction impeded the V2-mediated V1 protein redistribution and decrease TYLCV infection efficiency with delayed and mild symptoms. This report shows us a new explanation for the role of V2 in the nuclear export of V1 protein and TYLCV viral systemic infection.

## Introduction

Geminiviruses are a group of plant viruses with a circular, single-stranded DNA genome. Viruses in this family cause devastating diseases in crop plants, leading to agricultural losses worldwide [1–4]. While viral gene expression occurs in the cytoplasm, replication of geminiviruses occurs in the nucleus of infected host cells [5]. It is crucial that viral proteins involved in replication enter into the nucleus to execute their functions and in addition, newly synthesized viral genomic DNA are exported from the nucleus to the cytoplasm for further spread to adjacent cells and cause systemic infection through long-distance movement. Therefore, the nucleocytoplasmic shuttling of geminiviruses proteins and genomic DNA is of great significance for viral systemic infection and virus control.

Geminiviruses can be divided into two major groups based on their genomic components: one group is the monopartite geminiviruses, while the other group is the bipartite geminiviruses [5]. The bipartite geminiviruses genome is composed of two circular 2.5- to 2.8-kb ss-DNA molecules (DNA-A and DNA-B). The movement of bipartite geminiviruses requires two proteins, BV1 and BC1, that are encoded by DNA-B [6–10]. BV1 is a nuclear shuttle protein and plays an important role in the nucleocytoplasmic shuttling of viral genomic DNA [6–9, 11]. BC1 facilitates cell-to-cell movement after genomic DNAs are exported out of the nucleus.

The genome of monopartite geminiviruses contains only one component, DNA-A. Because monopartite geminiviruses lack the DNA-B component, the mechanism for movement of the virus is not clear yet. As DNA-A encode more viral proteins compared with DNA-B, and many of the proteins are multifunctional, which makes it more challenging to examine the nucleocytoplasmic shuttling of viral DNA of monopartite geminiviruses. Only a few viruses have been examined, such as *Maize streak virus* (MSV) and *Tomato yellow leaf curl virus* (TYLCV) [12, 13, 14]. It has been reported that V1 protein binds to viral genomic DNA and shuttles them between the nucleus and cytoplasm [11, 15, 16]. It was later reported that host proteins are also required for the process. Nuclear transporter KAP*α* helps TYLCV to enter the nucleus [17, 18] and exportin-*α* is required for the nuclear export of the C4 protein of *Tomato leaf curl Yunnan virus* (TLCYnV) [19]. In addition, nuclear shuttling of monopartite geminiviruses also involve viral proteins other than V1 protein, suggesting a protein complex may be involved [13, 19, 20]. However, it is unclear what viral proteins or how they work together to accomplish transportation between the nucleus and the cytoplasm.

TYLCV is a typical monopartite begomovirus in the family *Geminiviridae*. The single ssDNA genome has six open reading frames (ORFs) and an intergenic region (IR). Four ORFs (C1, C2, C3 and C4) are located on the complementary strand and the other two ORFs (V1 and V2) are located on the viral strand [21].

Replication-associated protein (Rep) encoded by C1, transcriptional activator protein (TrAP) encoded by C2 and replication enhancer protein (REn) encoded by C3 are all involved in viral replication. C4 protein is likely involved in symptom development and viral movement. V1 encodes the capsid protein (CP), which facilitates virion assembly and viral trafficking [1, 13, 22, 23]. V2 protein is a gene silencing suppressor at both the post-transcriptional stage (PTGS) [24] and the transcriptional level stage (TGS) [25]. V2 protein is also involved in the regulation of host defense responses [26] and viral movement [13], playing important roles in viral spread and systemic infection [27].

For the nucleocytoplasmic transportation of TYLCV, V1 protein is well-known as a nuclear shuttle protein and for its role in binding viral genomic DNA [13]. Early studies showed that V1 protein binds viral genomic ssDNA in the cytoplasm and moves them into the nucleus for replication [28, 29]. V1 protein also interacts with the host plant nuclear transporter protein to facilitate the entry of virus into the nucleus [17]. In addition, V1 protein also helps viral genomic DNA to move out of the nucleus for further translation and expression when offspring genomic DNA is synthesized [13]. This suggests that V1 protein is functionally equivalent to BV1 of bipartite geminiviruses. However, several lines of evidence suggest that other viral proteins, such as V2, are also involved [5, 13, 17, 30, 31, 32]. Rojas et al. found that the efficiency of nuclear export of viral DNA was enhanced by 20-30% in the presence of V2 protein, suggesting a role for V2 protein in the V1 protein-mediated nuclear export of viral genomic DNA [13]. However, the mechanism whereby V2 protein facilitates V1-mediated viral genomic DNA trafficking from the nucleus is unknown.

In this study, we demonstrate that V2 protein affects the subcellular localization of V1 protein by dramatically decreasing the nucleus-localized V1 protein in *Nicotiana benthamian* cells, possibly through host exportin-*α* (XPO I), which often mediates nuclear export of proteins. A specific interaction between V2 and V1 proteins has been identified by co-immunoprecipitation (Co-IP) and bimolecular fluorescence complementation (BiFC). Substitutions in cystine 85 of the V2 protein inhibits the V1-V2 interaction, blocks the effect of V2 protein on the subcellular localization of V1 protein, and causes delayed and mild symptom in plants. Our results indicate that V2 protein interacts with V1 protein, promotes the nuclear export of V1 protein, and plays an important role in viral systemic infection.

## Results

### V2 Protein Affects the Nuclear Localization of V1 Protein

TYLCV V1 protein is known as a nucleocytoplasmic shuttle protein that facilitates the transport of viral genomic DNA into and out of the nucleus. When expressed in cells of *Nicotiana benthamiana* by agroinfiltration as a YFP-tagged protein, V1-YFP, the YFP signal was found in both the nucleus and cytoplasm at 40 hours post infiltration (hpi) (Fig 1a), consistent with its role in nuclear transportation of viral genomic DNA.

**Fig 1.**
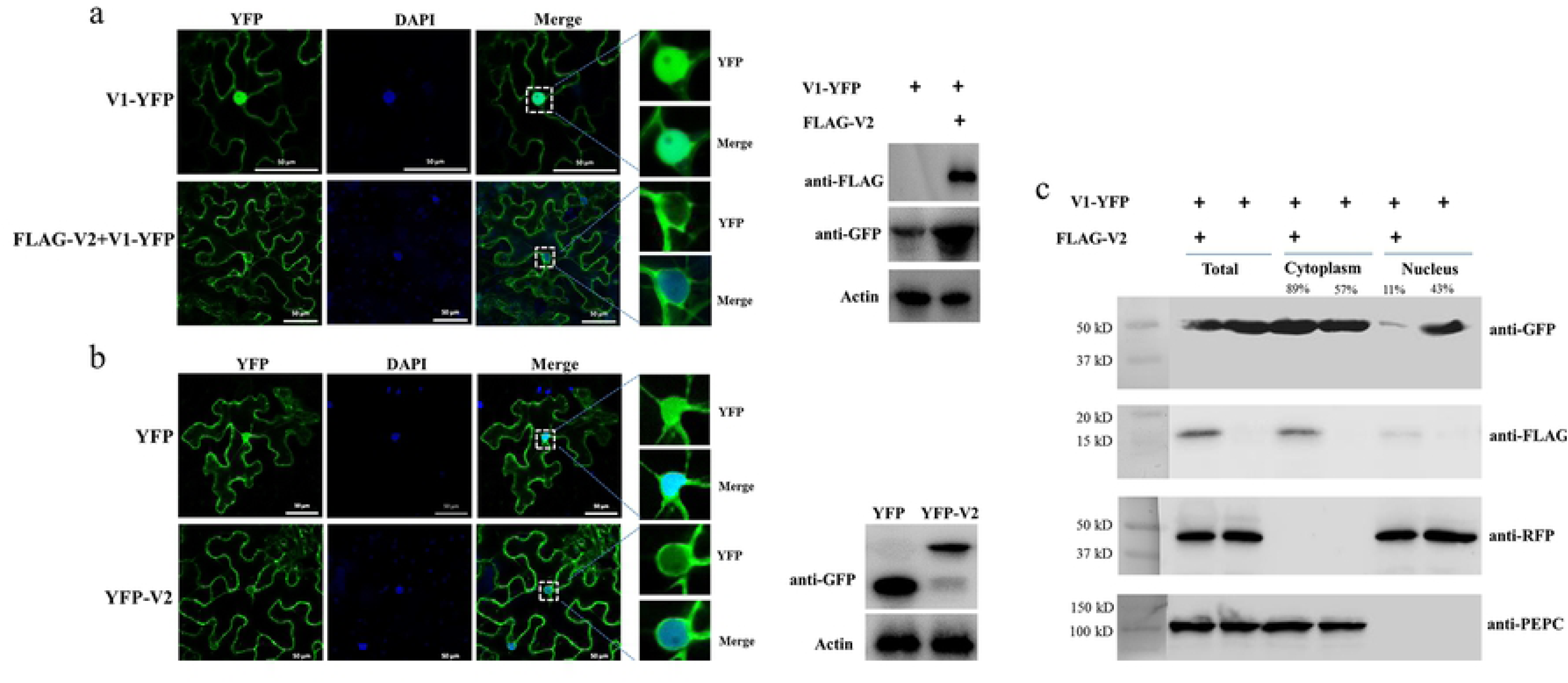
The effect of V2 protein on the nuclear distribution of V1 protein. (a) The localization of V1 protein in the absence or presence of V2 protein in *N. benthamiana* cells. V1-YFP was expressed in the absence or presence of FLAG-V2 and was detected either by confocal microscopy (left panel) or by Western blotting using an anti-GFP polyclonal antibody (right panel). DAPI stains DNA in the nucleus. Actin serves as a control for equal loading of total lysates. Bars: 50 µm. (b) The localization of V2 in *N. benthamiana* cells. The expressed YFP or YFP-V2 in epidermal cells of *N. benthamiana* leaves was detected either by confocal microscopy (left panel) or by Western blotting using an anti-GFP polyclonal antibody (right panel). DAPI stains DNA in the nucleus. Bars: 50 µm. (c) Nuclear-cytoplasmic fractionation assay of the distribution of V1 protein in the absence or presence of FLAG-V2 in H2B-RFP transgenic *N. benthamiana* plants. Nuclei were purified from plant tissues expressing V1-YFP in the absence or presence of FLAG-V2 using percoll density gradient centrifugation. Western blot analysis was conducted with antibodies specific to the indicated proteins using an anti-GFP polyclonal antibody or an anti-FLAG monoclonal antibody. PEPC was used as a marker for the cytoplasmic fraction and H2B-RFP was used as a marker for the nuclear fraction. Protein signal intensity was measured by using Adobe Photoshop CS6, with the cytoplasm plus the nucleus levels totaling as 100%.

Since V2 protein was reported to facilitate the export of viral genomic DNA from the nucleus [13], we tested whether V2 protein does so by promoting the nucleus export of the V1 protein. We first tested for the subcellular localization of V2 as a YFP-tag (V2-YFP) in *N. benthamiana* cells via agroinfiltration. The fluorescence signal was observed under a laser confocal microscope at 40 hpi. YFP-V2 was mainly present in the cytoplasm and perinuclear regions, but a much weaker signal was also present in the nucleus (Fig 1b). To further clarify the function of V2 protein in the nuclear export of TYLCV, we co-expressed FLAG-tagged V2 protein (FLAG-V2) with V1-YFP. Interestingly, only a weaker fluorescence signal of V1 protein was found in the nucleus compared to that of V1 protein alone (Fig 1a). To rule out the possibility that the weaker signal of V1-YFP in the nucleus was due to decreased expression and/or stability in the presence of V2, we checked the accumulation of V1-YFP by Western blotting. Our results showed that both V2 and V1 proteins were expressed well when co-expressed (Fig 1a). An increased accumulation of V1-YFP was sometimes noticed when it was co-expressed with FLAG-V2 than when it was expressed alone, indicating that the lower V1-YFP signal in the nucleus was not due to its decreased accumulation in the presence of FLAG-V2.

To confirm our visual observations, we performed a fractionation assay to separate the nucleus from the cytoplasm [19] and tested the localization of V1-YFP in the absence and presence of the V2 protein. To this end, we expressed FLAG-V2 and V1-YFP in Histone 2B (H2B)-RFP transgenic plants. As shown in Fig 1c, we only detected H2B-RFP in the nuclear fraction but not the cytoplasmic fraction; a cytoplasmic marker, phosphoenolpyruvate carboxylase (PEPC), was only present in the cytoplasm fraction, not in the nuclear fraction. Under such conditions, FLAG-V2 was primarily detected in the cytoplasm fraction but only weakly in the nucleus. Although V1-YFP was detected in both fractions when expressed alone, the amount in the nuclear fraction significantly decreased in the presence of FLAG-V2, which is consistent with the results based on fluorescence microscopy (Fig 1c). To provide a numeric reading, we set the sum of V1-YFP signal intensity in the cytoplasm and nucleus at 100%. In the absence of FLAG-V2, we found 43% of V1-YFP was associated with the nuclear fraction but decreased to 11% in the presence of FLAG-V2. We concluded from these results that V2 is able to change the nuclear localization of V1 protein.

### V2 Protein Interacts with V1 Protein

We set out to understand the underlying mechanism by which V2 protein affects the subcellular localization of V1 protein by first testing whether there is an interaction between V2 and V1 proteins by using a co-immunoprecipitation (Co-IP) assay. FLAG-tagged V2 (FLAG-V2) was co-expressed with YFP or V1-YFP in *N. benthamiana*. Total protein extracts were subject to immunoprecipitation by using FLAG-trap beads, and the resulting precipitates were analyzed using an anti-YFP antibody or an anti-FLAG antibody. Although a similar amount of FLAG-V2 was pulled down with FLAG-trap beads, only V1-YFP, but not YFP, was detected (Fig 2a), even though both YFP and V1-YFP were well expressed (Fig 2a).

**Fig 2.**
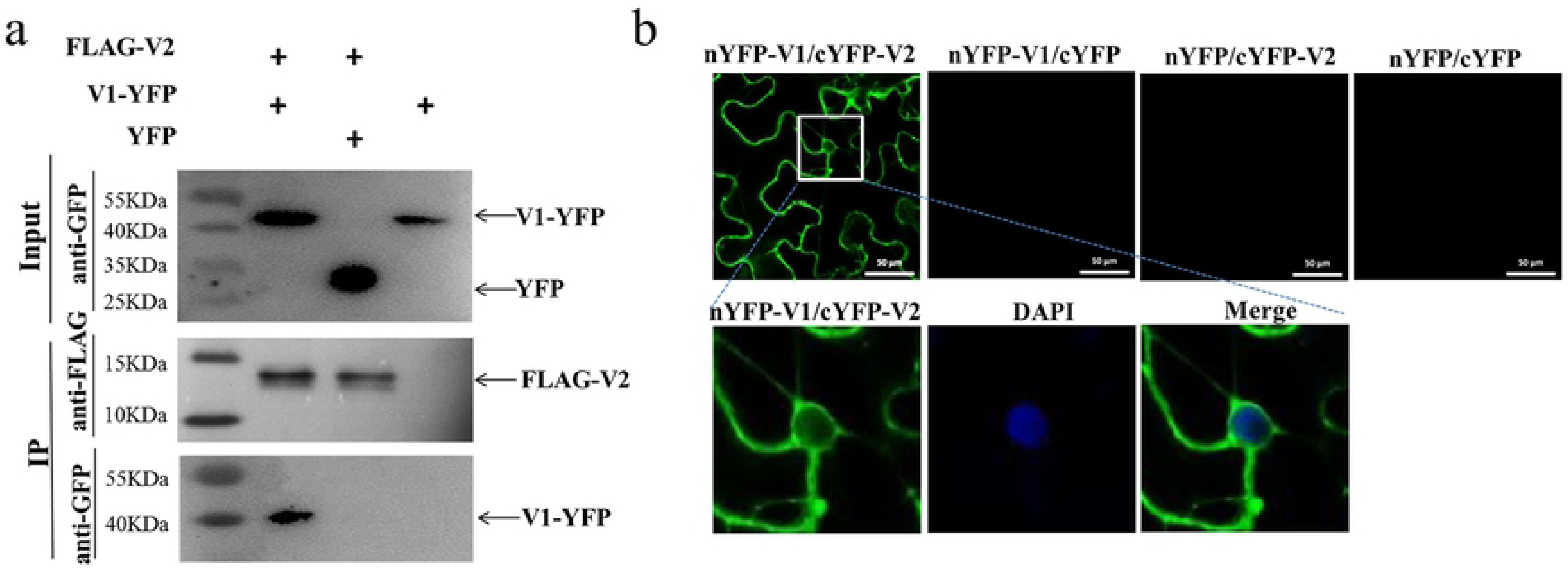
Identification of the interaction between V2 and V1 proteins. (a) Co-immunoprecipitation (Co-IP) analysis of the interaction between FLAG-V2 and V1-YFP. *N. benthamiana* leaves were co-infiltrated with *A. tumefaciens* cells harbouring expression vectors to express FLAG-V2 and V1-YFP (Lane 1), FLAG-V2 and YFP (Lane 2), or V1-YFP alone (Lane 3). Cell lysates were incubated with FLAG-trap beads (Sigma, USA). Samples before (Input) and after (IP) immunoprecipitation were analyzed by immunoblotting using anti-GFP or -FLAG antibody. (b) BiFC assays between V1 and V2 proteins in the leaves of *N. benthamiana*. Confocal imaging was performed at 48 hpi. V1 and V2 were fused to the N (nYFP) and C-terminal (cYFP) fragments of YFP, respectively. The V1-V2 interaction will lead to a reconstituted fluorescence signal. DAPI stains DNA in the nucleus. Bars: 50 µm.

The fact that V1 protein was co-precipitated with V2 protein suggests that V2 protein may bind to V1 to form a V1-V2 protein complex. To confirm the V1-V2 interaction and identify the location where the V1 and V2 proteins may form a complex, we used a bimolecular fluorescence complementation (BiFC) assay. A positive interaction between nYFP-V1 and cYFP-V2 was observed in both the cytoplasm and perinuclear region, as indicated by the presence of reconstituted green fluorescence (Fig 2b). We also noticed a faint fluorescence signal inside the nucleus. It should be noted that V1-YFP also localized in the cytoplasm and the perinuclear region when it was co-infiltrated with FLAG-V2 (Fig 1a), suggesting that V2 binds V1 protein at the perinucleus and the cytoplasm. No green fluorescence signal was generated when nYFP-V1 and cYFP, or nYFP and cYFP-V2, or nYFP and cYFP were co-expressed (Fig 2b), reinforcing a specific interaction between V2 and V1 proteins in plant cells.

### V2 Mediates the Nucleocytoplasmic Shuttling of V1 Protein Through Host Exportin-α

V2 can change the nuclear localization of the V1 protein, decreasing its accumulation in the nucleus. These results raised the possibility that V2 might help V1 protein export from the nucleus to the cytoplasm. Because the nuclear export of proteins is often mediated by exportin-α, we tested the subcellular localization of V2 upon treatment with leptomycin B (LMB), an inhibitor of exportin-α [33]. As expected, the level of nuclear-localized V2-YFP was increased after LMB treatment in epidermal cells of H2B-RFP transgenic *N. benthamiana* plants (Fig 3a), suggesting that V2 depends on exportin-α to move out of the nucleus. To confirm our observations, we performed a nuclear-cytoplasmic fractionation assay on H2B-RFP transgenic *N. benthamiana* plants expressing V2-YFP with or without LMB treatment. H2B-RFP and PEPC were used as nuclear- and cytoplasmic-localized marker proteins, respectively. About 32% of the total V2-YFP accumulated in the nucleus and increased to 54% with the LMB treatment (Fig 3b), agreeing well with our imaging results (Fig 3a).

**Fig 3.**
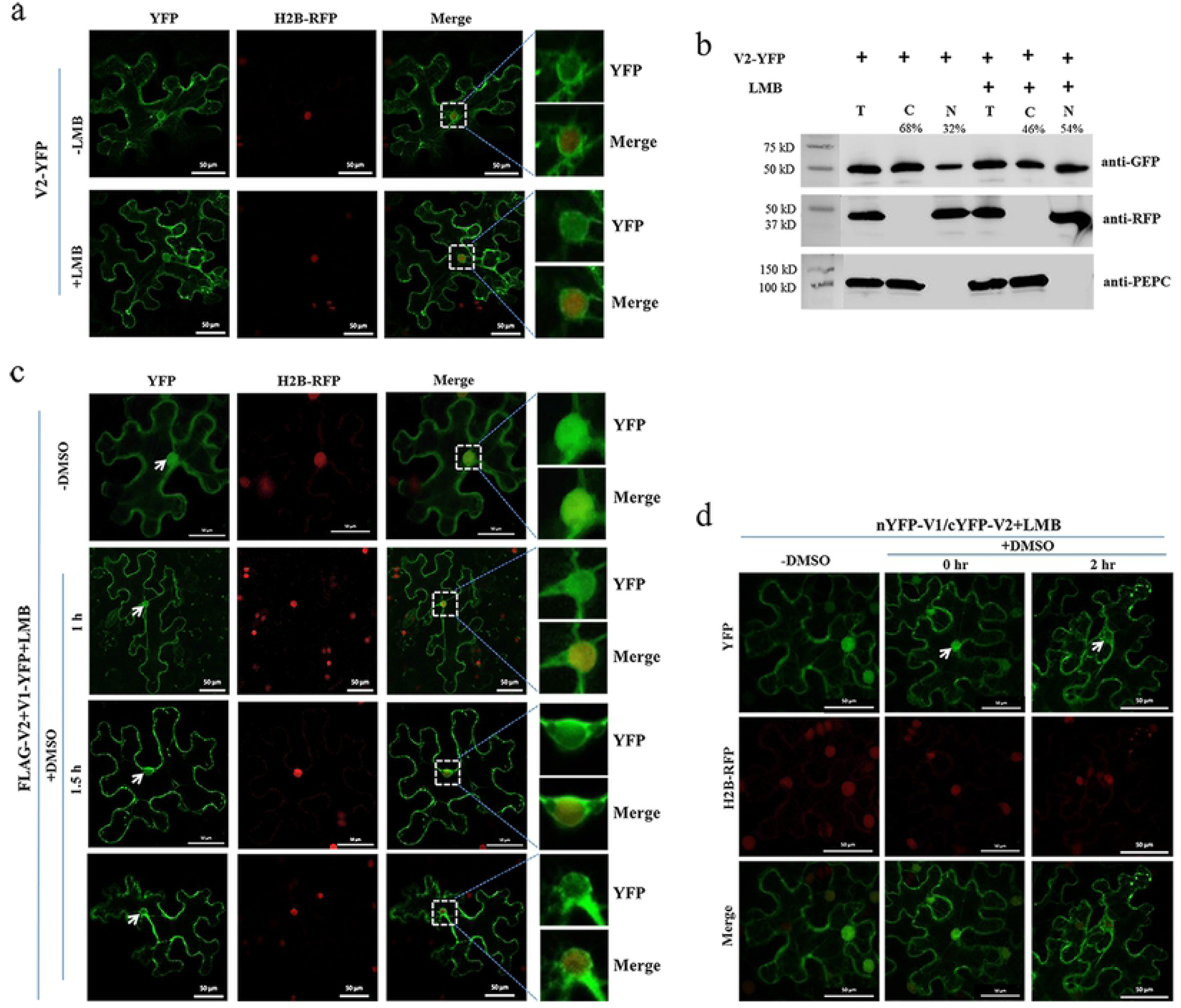
The V2-mediated nuclear export of V1 protein is dependent on exportin-α. (a) Subcellular distribution of V2-YFP without or with the LMB treatment in H2B-RFP transgenic *N. benthamiana* plants. Leaf tissues were first agroinfiltrated with V2-YFP for 40 hours and followed by 10 nM LMB for 2 hours. H2B-RFP signal represents the nucleus. Bars: 50 µm. (b) Nuclear-cytoplasmic fractionation analysis of the distribution of V2 with or without LMB treatment in H2B-RFP transgenic *N. benthamiana* cells. Western blot analysis was conducted with antibodies specific to the indicated proteins. PEPC was used as a marker for the cytoplasmic fraction and H2B-RFP as a marker for the nuclear fraction. Protein signal intensity was measured by using Adobe Photoshop CS6, the sum of cytoplasm plus the nucleus as 100%. (c) Subcellular distribution of V1-YFP co-expressed with FLAG-V2 upon the treatment of LMB and DMSO in H2B-RFP transgenic *N. benthamiana* plants. Leaf tissues expressing V2-YFP and FLAG-V2 were first infiltrated with 10 nM LMB for 2 hours followed by infiltration of 0.5% DMSO to degrade LMB. YFP signal was observed at specific time points as indicated. Arrows indicate the V1-YFP signal in or around the nucleus at different time points after the treatment with LMB and DMSO. H2B-RFP signal represents the nucleus. Bars: 50 µm. (d) Effects of the LMB treatment on the V1-V2 interaction as shown by BiFC in epidermal cells of H2B-RFP transgenic *N. benthamiana* plants. Plant tissues co-expressing nYFP-V1 with cYFP-V2 were treated with LMB for 2 h to inactivate exportin-α and then infiltrated with 0.5% DMSO to degrade LMB. Confocal micrographs were taken at different time points after DMSO treatment, as indicated. Arrow indicates localizations of the V1-V2 complex export from the nucleus. Nuclei of *N. benthamiana* leaf epidermal cells are indicated by the expression of the H2B-RFP transgene (red). Bars: 50 µm.

We also checked whether V2-mediated V1-YFP nuclear export can be affected by the LMB treatment. Co-expressed with FLAG-V2, V1-YFP had very low accumulation in the nucleus (Fig 1a), but a strong nuclear signal was observed after treatment with LMB (top panel, -DMSO, Fig 3c), suggesting that V2-mediated V1 protein nucleocytoplasmic shuttling is similar to the V2 protein export, which depends on exportin-α.

To confirm the specific effect of LMB on localizations of the V1 and V2 proteins, we further infiltrated LMB-treated cells with 0.5% dimethyl sulfoxide (DMSO), which degrades LMB. As expected, the V1-YFP signal was detected in the nucleus in the presence of FLAG-V2 and LMB at the beginning of DMSO treatment (top panel, -DMSO, Fig 3c). However, the V1-YFP signal in the nucleus decreased gradually after a longer DMSO treatment that eliminated the inhibitory effect of LMB (Fig 3c).

To verify the nucleocytoplasmic shuttling of the V1-V2 complex, we performed a BiFC assay applying the same treatments as above. In the presence of LMB only, the reconstituted YFP signal was strongly detected in the nucleus (Fig 3d), indicating that the V1-V2 complex was also present in the nucleus as well as in the cytoplasm and perinuclear region (Fig 2b). After DMSO treatment for 2 hours, the nucleus-localized YFP signal clearly diminished, indicating an exportin-α-mediated nucleocytoplasmic shuttling of the V1-V2 complex (Fig 3d). These results indicated that the nucleocytoplasmic shuttling of V1 protein is dependent on the V1-V2 interaction and exportin-α.

### The V2 Mutants C85A Abolishes the V1-V2 Interaction

To verify that the V1-V2 interaction plays a crucial role in the V1 nuclear export and to identify the approximate sites in V2 that are responsible for the interaction, we constructed six V2 mutants, each with single or double substitutions (Fig 4a). We then tested their interactions with V1 protein using the Co-IP assay. Among the six V2 mutants, only the V2^C85A^ mutant, which has a cysteine to alanine substitution in the residue at position 85, was not pulled down along with FLAG-V1 (Fig 4a,4b). Alanine substitution did not affect expression and stability of the V2^C85A^ mutant because V2-YFP and V2^C85A^-YFP accumulated at similar levels (top Input panel, Fig 4b).

**Fig 4.**
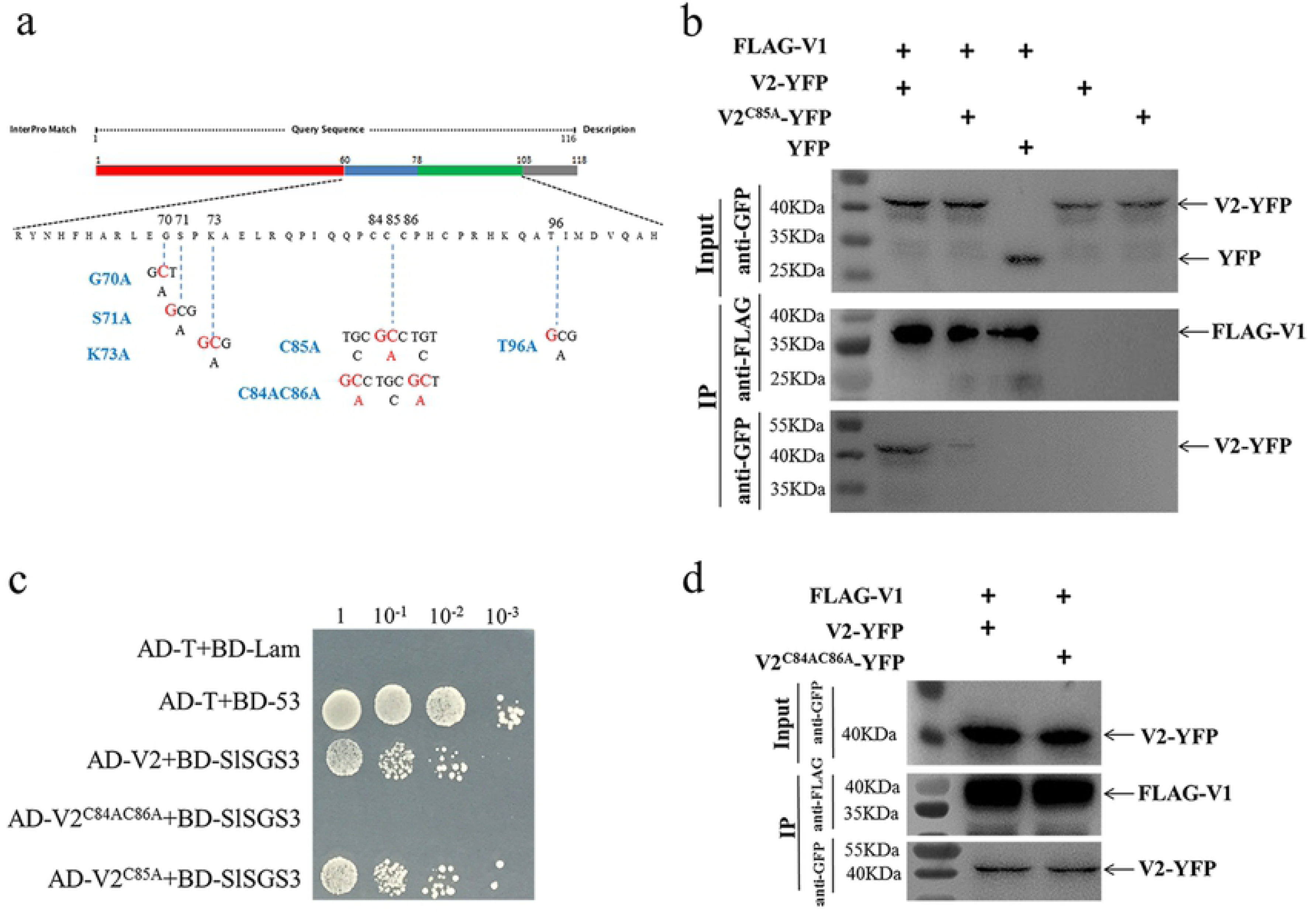
Identification of the critical sites in V2 protein that are responsible for the V1-V2 interaction. (a) Schematic illustration of the V2 protein. Nucleic acid and amino acids sequences of V2 mutants, V2^G70A^, V2^S71A^, V2^K73A^, V2^C85A^, V2^C84AC86A^, and V2^T96A^ are shown. (b) Co-IP assay to demonstrate the interaction between V1 and wt V2 or V2^C85A^. The Co-IP assay was performed as in Fig 2a. (c) Yeast-two hybrid (Y2H) detecting possible interactions between SlSGS3 and V2^C85A^ or V2^C84AC86A^. V2^C85A^ and V2^C84AC86A^ were fused with a GAL4 activation domain (AD-V2^C85A^ and AD-V2^C84AC86A^), and SlSGS3 was fused to a GAL4-binding domain (BD-SlSGS3), respectively. AH109 cells co-transformed with the indicated plasmids were subjected to 10-fold serial dilutions and plated on synthetic defined medium SD/-His/-Leu/-Trp medium to screen for positive interactions at 3 days after transformation. Yeast cells co-transformed with AD-T+BD-53 serves as a positive control; yeast cells co-transformed with AD-T+BD-Lam as a negative controls. (d) Co-IP assay to show the interaction between V1 and V2 or V2^C84AC86A^. The Co-IP assay was performed as in Fig 2a.

It is well-known that V2 protein is involved in PTGS by binding to tomato SGS3 (SlSGS3), a homologue of Arabidopsis SGS3 protein [34]. It has been confirmed that a mutant of V2 (V2 ^C84A/C86A^) does not interact with SlSGS3 and lost its function as a suppressor of gene silencing [34]. Given the fact that C85 is adjacent to C84 and C86, it is possible that V2^C85A^ may be dysfunctional not only in interacting with V1 protein but also with SlSGS3. To this end, we confirmed that V2^C85A^, but not V2^C84AC86A^, interacted with SlSGS3 in the yeast two-hybrid system (Fig 4c), indicating that the C85A substitution specifically blocked the V1-V2 interaction but did not disrupt other functions of the V2 protein, such as the ability to interact with SlSGS3 that leads to a block of host gene silencing-mediated host defense. To further confirm that C85, rather than C84 and C86, is required for different V2 roles, we also tested the ability of V2^C84A/C86A^ (Fig 4a) to interact with V1 protein. The Co-IP assay indicated that the V2^C84A/C86A^ mutant interacted with the V1 protein (Fig 4d). Taken together, the activities of the V2 protein in interacting with V1 protein and SlSGS3 can be separated, where the C85A mutation blocks V2 protein’s interaction with V1 protein but not with SlSGS3.

### The V2^C85A^ Mutant Fails to Redistribute the V1 Protein

After confirming that V2^C85A^ accumulated well and interacted with SlSGS3, we next checked the localization of V2^C85A^ by expressing YFP-tagged V2^C85A^ (V2^C85A^-YFP) in *N. benthamiana*. The fluorescence signal was observed in the cytoplasm and perinuclear region (Fig 5a), similar to wild-type (wt) V2-YFP, in 56% of cells expressing V2^C85A^-YFP (Fig 5b). In 44% of cells, however, the fluorescence signal was more spread than that of V2-YFP and was localized in an elongated region that does not exactly surround the DAPI-stained nucleus (Fig 5a). The nature of the localization remains to be determined. The C85A mutation did not affect the expression and stability of V2-YFP as V2^C85A^-YFP accumulated at a similar level as V2-YFP (Fig 5c). These results indicated that C85 has some effects on the perinuclear localization and the nuclear export function of the V2 protein.

**Fig 5.**
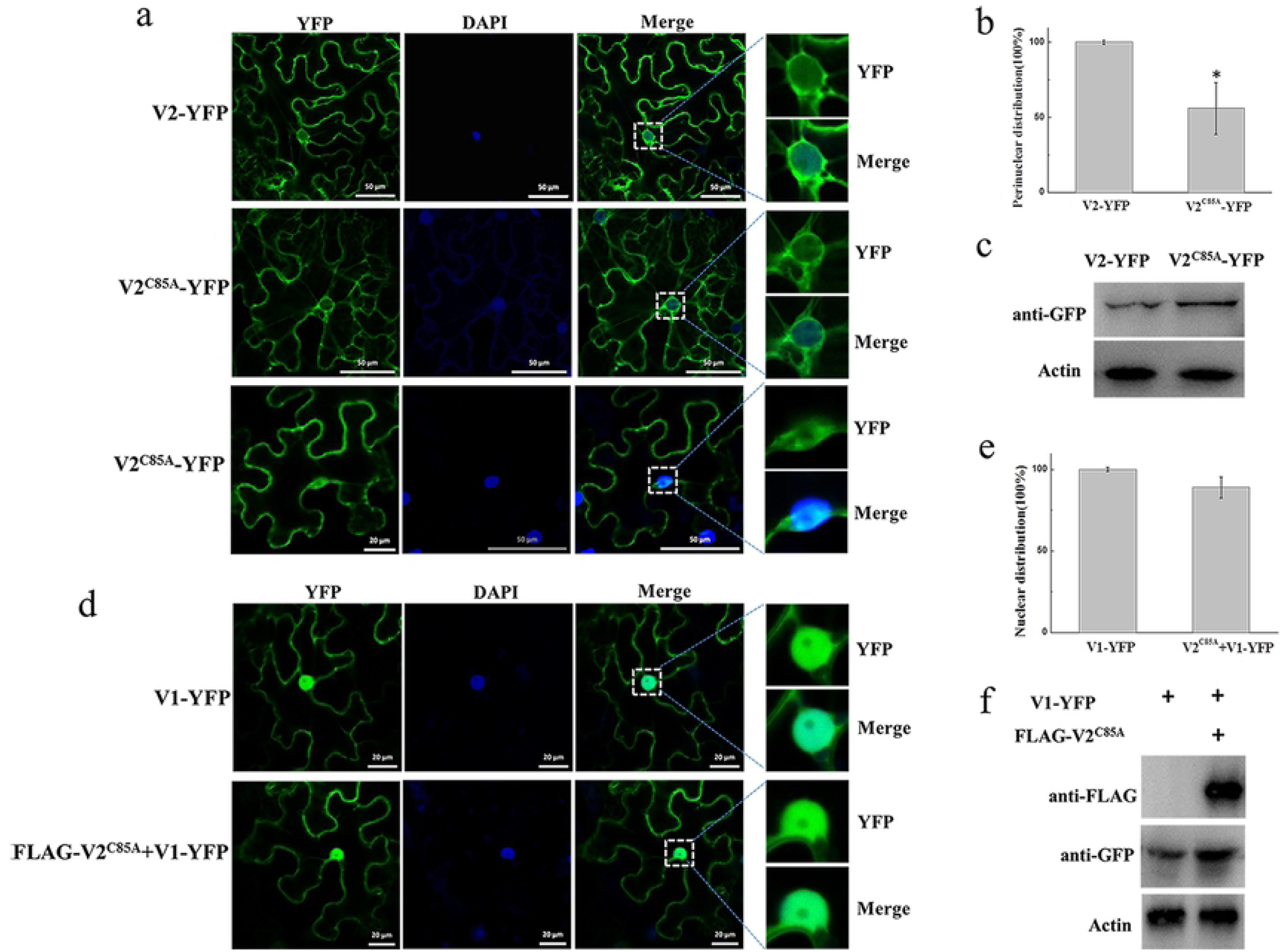
Characterization of the V2 ^C85A^ mutant. (a) Subcellular localization of V2 and V2^C85A^. DAPI stains DNA in the nucleus. Bars: 50 µm. (b) Quantification of perinuclear distribution of V2 and V2^C85A^. The number of cells with perinuclear distribution in different samples as in a. Experiments were repeated three times and 30 cells were observed in each trepeat. Values represent percentages of cells with a perinuclear distribution of YFP signal ± SD (standard deviation). Data were analyzed using Student’s t-test and asterisks denote significant differences between V2-YFP- and V2^C85A^-YFP-infiltrated leaves (*P < 0.05). (c) Western blot analysis showing accumulated V2 and V2^C85A^ using anti-GFP polyclonal antibody. Actin serves as a control for equal loading. (d) The localization of V1-YFP expressed alone or co-expressed with V2^C85A^ in *N. benthamiana* leaves. Bars: 20 µm. (e) Comparison of the nucleus-localized V1-YFP in the absence or presence of V2^C85A^. At least 150 cells were analyzed from three independent repeats (at least 50 cells from each experiment. Values represent mean ± SD relative to plants infiltrated with V1-YFP in the absence or presence of V2^C85A^. The data were analyzed using Student’s t-test and no significant difference was found for V1-YFP distribution between V1-YFP and V1-YFP+ FLAG-V2^C85A^. (f) The accumulated V1-YFP and FLAG-V2^C85A^ as shown by Western blot analysis.

To test the effect of V2^C85A^ on the localization of V1, FLAG-V2^C85A^ was co-expressed with V1-YFP in *N. benthamiana* cells. A strong V1-YFP signal was detected in the nucleus in the presence of FLAG-V2^C85A^, similar to that when V1-YFP was expressed alone (Fig 5d). Among 50 cells that were observed for the localization of V1 protein, no obvious difference in the V1-YFP distribution pattern was observed in the absence or presence of V2^C85A^ (Fig 5e), suggesting that V2^C85A^ was not able to affect the nuclear localization of V1 protein. Because V1 protein accumulated at similar levels in the absence or presence of V2^C85A^ (Fig 5f), we propose that the disrupted V1-V2 interaction is responsible for the failed redistribution of V1-YFP in the presence of V2^C85A^.

### A C85 Substitution in V2 Protein Delays Viral Systemic Infection

To assess the role of the V1-V2 interaction in viral infection, we constructed an infectious TYLCV clone with a substitution in the C85 of V2 coding sequence. As V2 ORF overlaps with V1 ORF in the TYLCV genome, mutations in V2 may affect V1 amino acid sequence. To ensure that changes in C85 has no effect on V1 protein in TYLCV genome, the C85S mutation was introduced into a TYLCV clone to generate TYLCV-C85S, based on the fact that V2-C85S mutant did not interact with V1 protein (S1a Fig), but interacted with SlSGS3 (S1b Fig), and did not affect the subcellular localization of V1-YFP (S1c Fig). TYLCV-C85S and TYLCV were used to inoculate *solanum lycopersicum* and *N. benthamiana* plants.

Fifteen tomato plants were inoculated with either wt TYLCV or TYLCV-C85S. Typical symptoms such as chlorosis on leaves were first observed at 16 days post agro-infection (dpai) on tomato plants inoculated with TYLCV. No obvious symptoms were observed in the plants inoculated with TYLCV-C85S at 16 dpai (Fig 6a). The vast majority of TYLCV-C85S-inoculated tomato plants remained symptomless even at 32 dpai and only 1-2 plants among 15 eventually developed mild symptoms eventually, such as leaf yellowing (Fig 6b). Real-time PCR showed that viral DNA accumulation was much lower in plants inoculated with TYLCV-C85S than in plants inoculated with wt TYLCV (Fig 6c). Almost no virus particles accumulated in the TYLCV-C85S-inoculated plants based on a Western blotting assay using an anti-CP antiserum at 16 dpai (Fig 6d).

**Fig 6.**
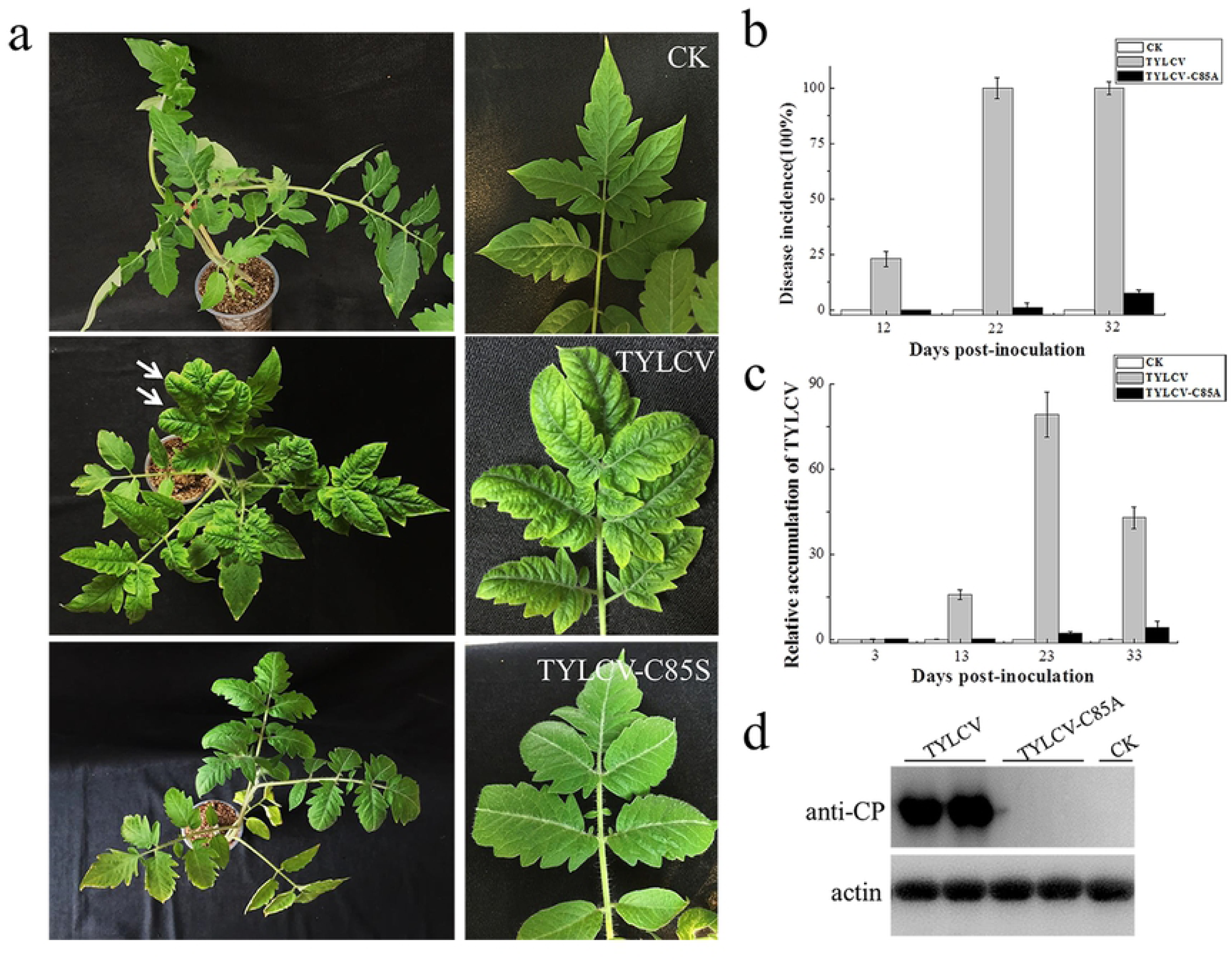
Effects of the C85S mutation on viral infection and viral accumulation in TYLCV-inoculated tomato plants. (a) Symptoms in plants that were agro-inoculated with wt TYLCV or TYLCV-C85S at 16 dpai. CK represents mock-inoculated plants. Arrows point to the yellowing and curly leaves. (b) The infection time course of wt TYLCV or TYLCV-C85S infection. Values represent percentages of systemically infected plants at different DPAI and are given as mean ± SD of triplicate experiments. In each experiment, 15 plants were inoculated and three independent repeats were performed to confirm the results. (c) Viral DNA levels in plants as measured by RT-PCR. Plants were inoculated with a wt TYLCV infectious clone or the TYLCV-C85S, or mock-inoculated (CK). Accumulated levels of *V1 (CP)* were tested at 3, 13, 23, and 33 dpai. Tomato leaves were agroinfiltrated with CK, TYLCV, or TYLCV-C85S. Total RNAs were extracted from newly emerged leaves. Values represent the mean relative to the CK-treated plants (n=3 biological replicates) and were normalized with *SlActin* as an internal reference. (d) Western blot analyses of accumulated V1 (CP), which represents viral particles, in CK, TYLCV and TYLCV-C85S inoculated plants at 16 dpai.

Similar results were also obtained in TYLCV-C85-inoculated *N. benthamiana* plants. All wt TYLCV-inoculated plants showed typical symptoms at 22 dpai, such as leaf yellowing and curling, but only one out of fifteen plants inoculated with TYLCV-C85S showed mild symptoms (Fig 7a, 7b). Accumulated TYLCV genomic DNA (Fig 7c) and virus particles (Fig 7d) in systemic leaves of TYLCV-C85S-inoculated plants were much lower than those in wt TYLCV-inoculated plants.

**Fig 7.**
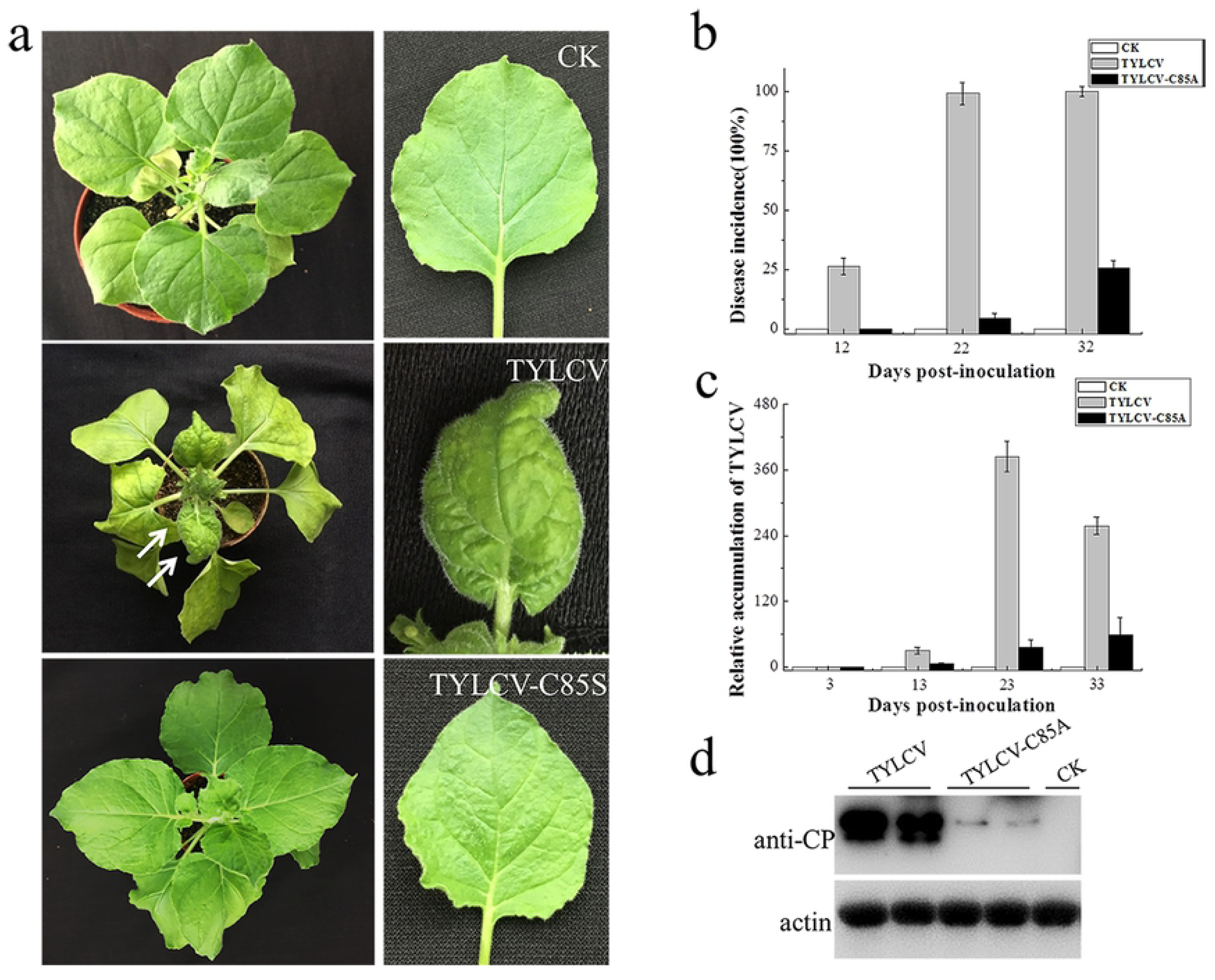
Effects of the C85S mutation on viral infection and viral accumulation in TYLCV-inoculated *N. benthamiana*. (a) Symptoms in plants that were agro-inoculated with wt TYLCV or TYLCV-C85S at 16 dpai. CK represents mock-inoculated plants. Arrows point to the curly leaves. (b) The infection time course of wt TYLCV or TYLCV-C85S infection. Values represent percentages of systemically infected plants at different DPAI and are given as the mean ± SD of triplicate experiments. In each experiment, 15 plants were inoculated and three independent experiments were performed to confirm the results. (c) Viral DNA levels in plants inoculated with wt TYLCV infectious clone or TYLCV-C85S. Accumulated levels of *V1 (CP)* were tested at 3, 13, 23, and 33 dpai. *N. benthamiana* leaves were agroinfiltrated with CK, TYLCV, or TYLCV-C85S. Total RNAs was extracted from newly emerged leaves. Values represent the mean relative to the CK-treated plants (n=3 biological replicates) and were normalized with *NbActin* as an internal reference. (d) Western blot analyses of accumulated V1 (CP), which represents viral particles, in CK, TYLCV and TYLCV-C85S inoculated plants at 16 dpai.

These results collectively showed that the mutation at C85 of V2 protein caused significant low levels of virus accumulation in the systemic leaves and dramatic decrease of the infection efficiency with delayed and mild symptoms.

## Discussion

Because genome replication of geminiviruses takes place in the nucleus of the infected host cells [5], it is crucial to transport the viral offspring DNAs from the nucleus back to the cytoplasm for intracellular, cell-to-cell, and long-distance movement. In bipartite geminiviruses, it is well-known that BV1 protein encoded by the DNA-B component facilitates trafficking of the viral genome into and out of the host nucleus [6–9, 35]. However, monopartite geminiviruses, which do not contain the DNA-B component, does not encode BV1 protein. So, the viral DNA shuttling between the nucleus and the cytoplasm is accomplished by protein or a protein complex encoded by the DNA-A component. It has been reported that V1 protein of monopartite geminiviruses mediates the import and export of viral DNA [13, 17, 28].

However, V1 protein might not be the only viral protein that is involved in the nucleocytoplasmic shuttling of TYLCV. Previous reports based on triple microinjection experiments revealed that the nuclear export of DNA was enhanced 20–30% in the presence of V2 (V2+V1+viral DNA), suggesting that V2 enhances nuclear export of viral DNA [13]. But the mechanism by which V2 protein promotes viral DNA export is unclear. We report here that V2 may facilitate viral DNA export by interacting with V1 and promoting the nuclear export of V1 protein.

In this study, we found that V2 protein localized primarily in the perinuclear region and the cytoplasm (Fig 1b). A very weak signal was also present in the nucleus (Fig 1b, 1c), but upon treatment with the exportin-α inhibitor LMB, the amount of nucleus-localized V2 protein increased significantly (Fig 3a), suggesting V2 protein shuttles between the nucleus and the cytoplasm but is quickly exported out of the nucleus via exportin-α. It is unclear, however, how V2 imported into the nucleus.

Our work indicates that V2 plays a critical role in the nuclear export of V1 protein as the nucleus-localized V1 diminished when V2 was present (Fig 1a). Supporting this notion, we found that LMB treatment, which prevented V2 from exporting out of the nucleus (Fig 3a), blocked V2-mediated nuclear export of V1 (Fig 3c). We also showed that the specific V1-V2 interaction is closely related to V1 trafficking. The V1-V2 interaction primarily occurred at the perinuclear region and the cytoplasm (Fig 2b) but was strongly detected in the nucleus upon LMB treatment (Fig 3d), suggesting that they may be in a complex or complexes throughout the viral replication and movement in infected cells. In addition, it also suggested that LMB only specifically blocked V2’s transport out of the nucleus but had no effect on the V1-V2 interaction. However, our data are not able to determine whether V2 mediates the nuclear import of V1 protein. In addition, our results do not rule out the possibility that other viral proteins, such as C4 protein, may also be involved in this process.

Cysteine at 85 of V2 was found to be crucial for the V1-V2 interaction because substitutions of Cys85 with alanine (Fig 4b) or serine (S1a Fig) led to substantially inhibited interaction with V1 and thus, its ability to facilitate V1’s transport out of the nucleus (Fig 5d, 5e for C85A and S1c Fig for C85S). Because V1 is known for binding to and facilitating nucleocytoplasmic trafficking of viral DNA [13, 17, 28], and because V2 facilitates the nuclear export of viral DNA along with V1 [13], we propose that the V2-promoted nuclear export of viral DNA is via the V1-V2 interaction. Our hypothesis is consistent with our results that the TYLCV-C85S mutant, which has the C85S mutation incorporated into an infectious TYLCV clone, led to the delayed onset of symptoms with only mild symptoms only in 1-2 tomato or *N. benthamiana* plants out of a total of 15 (Fig 6a, 6b, 7a, 7b). Real-time PCR and Western blotting assay results confirmed significant reduction in viral accumulation in TYLCV-C85S-inoculated plants compared with those plants inoculated by wt TYLCV (Fig 6c, 6d, 7c, 7d). These results showed that the cysteine at 85 of V2 was very important for viral systemic infection. The mutant on C85 caused V2 to lose its ability to bind with V1 and lead to V2 being unable to help with V1 accumulation at the perinuclear region nor participate in V1-mediated nuclear export of viral genomic DNA, which eventually affected the viral systemic infection.

In monopartite geminiviruses, V2 is a multifunctional protein that is involved in suppressing host PTGS and TGS, pathogenicity and systemic infection [5,30,31,32]. Substitution in cysteine 85 may affect functions other than its interaction with V1, especially since both Cys84 and Cys86 are critical for interacting with SlSGS3 and the suppression of gene silencing [34]. We found that even though the C85A (Fig 4b) and C85S (S1a Fig) mutants failed to interact with V1 protein and thus, V1’s trafficking out of the nucleus (Fig 5d, 5e, S1c), but both interacted with SlSGS3 (Fig 4c, S1b), suggesting that C85A and C85S mutants maintain their activity as gene silencing suppressors. These results also are consistent with the notion that the C85S mutation delays viral systemic infection by affecting V1-mediated viral genomic DNA transportation from the nucleus to the cytoplasm, not by disturbing gene silencing-mediated host defense. However, we cannot totally rule out the possibility that other V2-mediated viral infection step(s) besides viral DNA trafficking are affected by the C85S mutation.

Our data also showed that the C84A/C86A double mutant interacted with V1 (Fig 4d) but not SlSGS3 (Fig 4c), indicating that C84 and C86 are not related to V2’s ability to interact with V1. Our results therefore revealed that motifs responsible for V1’s nuclear export and gene silencing suppressor activity are independent from one another.

Our results indicate that V2 binds to V1 protein and facilitate the nuclear export of V1. During TYLCV infection, V1 mediates both nuclear import and export of viral DNA. The equilibrium between nuclear targeting and nuclear egress is changed upon completion of replication and the V1-V2 interaction can improve the nuclear export of the V1-DNA complex. Thus, viral DNA will be preferentially transported out of the nucleus for subsequent infection events. In the presence of the V2^C85S^ mutant, the nuclear export of V1 is slowed down or eliminated and therefore, viral DNA and subsequent viral cell-to-cell and systemic movement is delayed. However, we cannot totally rule out that the V1-V2 complex is also required for intracellular, cell-to-cell, and/or long-distance movement besides nuclear export of V1 protein and V1-mediated viral offspring DNAs.

Based on our findings here, we propose a working model for the role of V2 in V1-mediated nuclear export of TYLCV genomic DNA (Fig 8). When offspring viral genomic DNA are produced in the nucleus, they are bound by V1 [29]. A V2-V1-viral DNA complex is subsequently formed via a specific interaction between V1 and V2 and, with the help of exportin**-**α, V2 facilitates the V1-viral DNA complexes to egress from the nucleus to the perinucleus and the cytoplasm with an enhanced efficiency. Eventually TYLCV then spreads to adjacent cells and upper leaves, which results in a systemic infection. The infection efficiency and the accumulation of TYLCV in the systemic leaves are dramatically decreased without a defective V2-V1 interaction.

**Fig 8.**
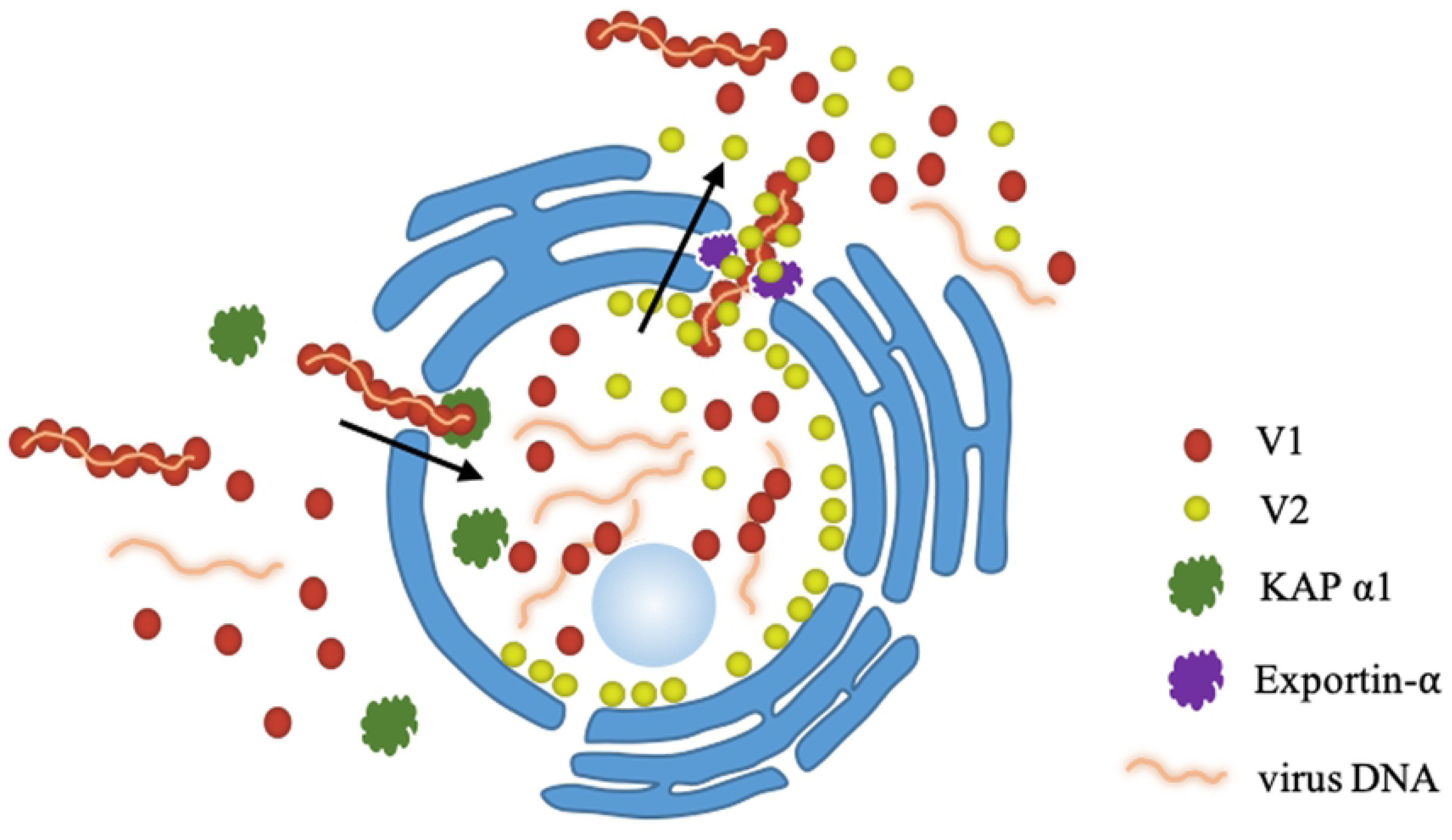
A working model proposed for V2-mediated nucleocytoplasmic trafficking of V1 protein. Viral genomic DNAs are bound by V1 and import into the nucleus with the help of KAP α1 [17, 18]. Via the specific interaction between V1 and V2, a V2-V1-ssDNA complex is formed. With the help of exportin**-**α, V2 facilitates the V1-ssDNA complexes to exit the nucleus to the perinucleus and the cytoplasm.

In summary, our results revealed that one mechanism of V2 protein’s involvement in viral DNA transportation is to promote V1-mediated viral transportation from the nucleus to the perinuclear region and the cytoplasm, which is with a specific interaction with V1, form V2-V1-viral DNA complex, and via host exportin-α. However, whether V2 promotes the ability of V1 to bind viral DNA and whether the V1-V2 interaction works after nuclear transportation require further research.

## Materials and Methods

### Plant Materials and Growth Conditions

Transgenic *Nicotiana benthamiana* plants expressing a nuclear marker, H2B-RFP (red fluorescent protein fused to the C terminus of histone 2B) [36], were kindly provided by Dr. Xiaorong Tao (Nanjing Agricultural University, Nanjing, China).

All agro-infiltration experiments were performed in wild-type (wt) or H2B-RFP transgenic *N. benthamiana*. Plants were grown in a growth chamber (ModelGXZ500D, Jiangnan Motor Factory, Ningbo, China) at 26°C (16 h, light) and 22°C (8 h, dark) for 4-6 weeks before being infiltrated with the agrobacterium. After infiltration, the plants were kept under the same growth conditions.

### Plasmid Construction

The coding sequences of TYLCV V2 and V1 genes were amplified from the cDNA of a TYLCV-infected tomato plant from Jiangsu Province, China (GenBank accession number GU111505) [37], using corresponding primers (S1 Table). Site-specific mutants of V2^G70A^, V2^S71A^, V2^K73A^, V2^C85A^, V2^C84AC86A^, V2^C85S^, V2^T96A^ were synthesized (Invitrogen, China) and confirmed by sequencing (Fig 4a).

To investigate the subcellular localization, the TYLCV V2 (*Bgl*II), V1 genes (*Bam*HI) and V2^C85A^, V2^C84AC86A^, V2^C85S^ were amplified using specific primers (Supplementary Table S1). Yellow fluorescent protein (YFP) tag was inserted between the CaMV 35S promoter and the 35S terminator (35St) in the pCambia1300 binary vector to construct the p1300-YFP vector as previously described [38]. Then, amplified products were individually inserted either into the *Bam*HI or *Bgl*II (compatible with *Bam*HI) site of the p1300-YFP vector to fuse in frame with YFP at the N-terminus or C-terminus to generate V2-YFP, V1-YFP, YFP-V2 and V2^C85A^-YFP, V2^C84AC86A^-YFP, V2^C85S^-YFP.

To make BiFC vectors, full-length coding sequences of V2 and V1 genes were amplified using the primers listed in Supplemental Table S1, then V1 was cloned into the *Bam*HI site as a fusion with the N-terminal fragment of YFP and V2 was cloned into the *Bam*HI site as a fusion with the C-terminal fragment of YFP, resulting in nYFP-V1 and cYFP-V2.

FLAG tagged V2 and V1 were amplified by PCR using specific primers (S1 Table) and inserted into the *Bam*HI site between the 35S promoter and the 35St in the pCambia1300 binary vector to generate FLAG-V2 and FLAG-V1 for the further Co-IP experiments.

For the yeast two-hybrid assay, V2, V2^C85A^, V2^C84AC86A^, V2^C85S^ were amplified and inserted into the *Nde*I/*Eco*RI-digested pGADT7 vector and SlSGS3 was cloned into the *Nde*I/*Bam*HI-digested pGBDT7 vector.

### Agro-infiltration Assays in *N. benthamiana*

Target vectors were transformed into *A. tumefaciens* strain GV3101 by electroporation. Agrobacterial cultures were harvested when the OD600 was approximately 0.8–1.0, collected by centrifugation, resuspended in the induction buffer (10 mM MgSO4, 100 mM 2-N-morpholino ethanesulfonic acid [pH 5.7], 2 mM acetosyringone), and incubated for 2 h at room temperature. The suspensions were then adjusted to OD600=0.5 and infiltrated into 4- to 6-week-old wt *N. benthamiana* or H2B-RFP transgenic *N. benthamiana* leaves for the further experiments.

### Subcellular Localization of Proteins

P1300-YFP, YFP-V2, V2-YFP, V1-YFP, V2^C84AC86A^ and V2^C85A^-YFP were individually introduced into *A. tumefaciens* strain GV3101 through electroporation. Leaves of 4-week-old *N. benthamiana* plants were infiltrated with *A. tumefaciens* harbouring the designated constructs. At 40 hours post infiltration (hpi), plants leaves were excised and YFP fluorescence was examined in epidermal cells using confocal microscopy (Zeiss LSM 880). The microscope was configured with a 458–515 nm dichroic mirror for dual excitation and a 488-nm beam splitter to help separate YFP fluorescence.

### Bimolecular Fluorescence Complementation (BiFC) Assay

BiFC experiments were performed as previously described [39] with minor modifications. nYFP-V1 and cYFP-V2 were introduced individually into *A. tumefaciens* strain GV3101 by electroporation. After overnight growth and activation, agrobacterium cultures were combined and infiltrated into leaves of *N. benthamiana* as above. After agroinfiltration, *N. benthamiana* plants were grown in a growth chamber with a 16 h light/8 h dark cycle. YFP fluorescence was observed and photographed by using confocal microscopy (Zeiss LSM 710) at 48 hpi. YFP was observed under a mercury lamp light using a 488-nm excitation filter. Photographic images were prepared using ZEN 2011SP1.

### Co-Immunoprecipitation

The Co-IP assay was performed as previously described [38]. 40 h after infiltration, *N. benthamiana* leaves were harvested and ground in liquid nitrogen. Proteins were extracted in IP buffer (40 mM Tris-HCl at pH 7.5, 100 mM NaCl, 5 mM MgCl2, 2 mM EDTA, 2×EDTA-free proteinase inhibitor, 1 mM PMSF, 4 mM DTT, 1% glycerol, and 0.5% Triton-X100). After centrifugation, the supernatant was mixed with FLAG-conjugated beads (Sigma, USA). After 1 h incubation at 4°C, the beads were washed six times with IP buffer, resuspended in 2×SDS gel loading buffer, and boiled for 10 min. The samples were loaded onto a 12% (vol/vol) SDS/PAGE gel and target proteins were detected using a polyclonal anti-GFP antibody (GenScript, USA) or a monoclonal anti-FLAG (Sigma, USA) antibody.

### Yeast Two-Hybrid Assay

The yeast two-hybrid system was used to examine interactions between V2, V2^C85A^, V2^C84AC86A^ and SlSGS3. V2, V2^C85A^, and V2^C84AC86A^ were cloned into the activation domain (AD)-containing vector and SlSGS3 was cloned into the vector harboring the DNA binding domain (BD). Both constructs were transformed into *Saccharomyces cerevisiae* strain AH109. The plasmid pairs BD-53 and AD-recT served as a positive controls, while the plasmid pairs BD-Lam and AD-recT were used as a negative controls. Transformants were grown at 30°C for 72 h on SD/-His/-Leu/-Trp/ synthetic medium to test for protein-protein interaction.

### Nuclear-Cytoplasmic Fractionation Assay

Nuclear-cytoplasmic fractionation assays were performed as described previously [40] with minor modifications. Infiltrated leaves were harvested and mixed with 2 mL/g of lysis buffer (20 mM Tris-HCl, pH 7.5, 20 mM KCl, 2 mM EDTA, 2.5 mM MgCl2, 25% glycerol, 250 mM Sucrose, 5 mM DTT and 10 mM protease inhibitor). The homogenate was filtered through a double layer of Miracloth. The supernatant, consisting of the cytoplasmic fraction, was centrifuged at 10,000 rpm for 10 min at 4°C and collected. The pellet was resuspended with 500 µL of NRB2 (20 mM Tris-HCl, pH 7.5, 0.25 M Sucrose, 10 mM MgCl2, 0.5% Triton X-100, 5 mM b-mercaptoethanol and 10 mM protease inhibitor) and overlaid on top of 500 µL NRB3 (20 mM Tris-HCl, pH 7.5, 1.7 M Sucrose, 10 mM MgCl2, 0.5% Triton X-100, 5 mM b-mercaptoethanol and 10 mM protease inhibitor). These were centrifuged at 12,000 rpm for 40 min at 4°C. The final nuclear pellet was resuspended in 400 µL lysis buffer. As quality controls for the fractionation assays, PEPC protein and H2B-RFP were used as a cytoplasmic and a nuclear marker, respectively.

### Leptomycin B Treatment Assays

Leptomycin B (LMB) Treatment Assays were performed as previously described [19] with minor modifications. LMB (Fisher Scientific, USA) was dissolved in ethanol to prepare 10 mM stock solutions. For *in vivo* treatment of *N. benthamiana* leaves, stock solutions were diluted in water to prepare solutions of 10 nM LMB. Agroinfiltrated *N. benthamiana* leaves expressing the protein of interest at 40 hpi were infiltrated with 10 nM LMB. 2 h after LMB treatment, the leaves were cut and mounted on a glass slide for confocal imaging. When needed, DMSO was further infiltrated into LMB-treated leaves and tissues were harvested at the specified time points, namely 1, 1.5, and 2 h.

### TYLCV constructs for Agrobacterium-mediated inoculation

For the construction of infectious clones of TYLCV containing V2^C85S^, a full length TYLCV mutant, TYLCV-C85S (the cysteine residue on V2 at amino acid 85 was changed to serine), was synthesized (Invitrogen, China). Then the full DNA-A of TYLCV-C85S was amplified using the primers listed in S1 Table and inserted into the pGEM-T Easy (Promega, USA) vector to produce pGEM-1A-C85S. After sequence confirmation, a 2183 nucleotide (nt) fragment was excised from pGEM-1A with *Bam*HI and *Sac*I, then subcloned into the *Bam*HI-*Sac*I sites of the binary vector pBinPLUS to produce pBinPLUS-0.8A. The full-length of TYLCV-C85S was digested from pGEMT-1A with *Bam*HI and inserted into pBinPLUS-0.8A at its unique *Bam*HI site to get pBinPLUS-1.8A, the infectious clone of TYLCV-C85S. The wt TYLCV infectious clone was constructed as previously described [38].

Agrobacterium cultures harboring TYLCV constructs were injected into the stem of *S. lycopersicum* and *N. benthamiana* with a syringe. Inoculated plants were grown in an insect-free cabinet with supplementary lighting corresponding to a 16-h day length.

### RNA extraction and qRT-PCR analysis

Total RNA was extracted from mock (Agrobacterium-carrying empty vector)-, wt TYLCV- or TYLCV-C85S-infiltrated *S. lycopersicum* or *N. benthamiana* leaves at different time periods using the Trizol Reagents (Life technologies, USA) and treated with DNase I following the manufacturer’s instructions (PrimeScript RT reagent Kit with gDNA Erase, Takara, Japan). cDNA synthesized from reverse transcription of RNA samples was used to determine the mRNA expression levels of target genes as well as for quantifying TYLCV accumulation levels at the date indicated. *SlActin* or *NbActin* was used as an internal control for tomato or *N. benthamiana*, respectively.

## Acknowledgements

We thank Dr. Sue Tolin (Virginia Tech, USA) and Dr. Janet Webster (Virginia Tech, USA) for critical reading of the manuscript. We are grateful to Xiaorong Tao (Nanjing Agricultural University, China) for kindly providing the H2B-RFP transgenic *N. benthamiana*.

## Author contributions

Y.Z., Y.J., X.W. and W.Z. designed the project. W.Z., S.W. and E.B. conducted experiments. All authors analyzed the data and reviewed the manuscript. W.Z., Y.J. and X.W. wrote the paper.

## Supporting Information

**S1 Table.**
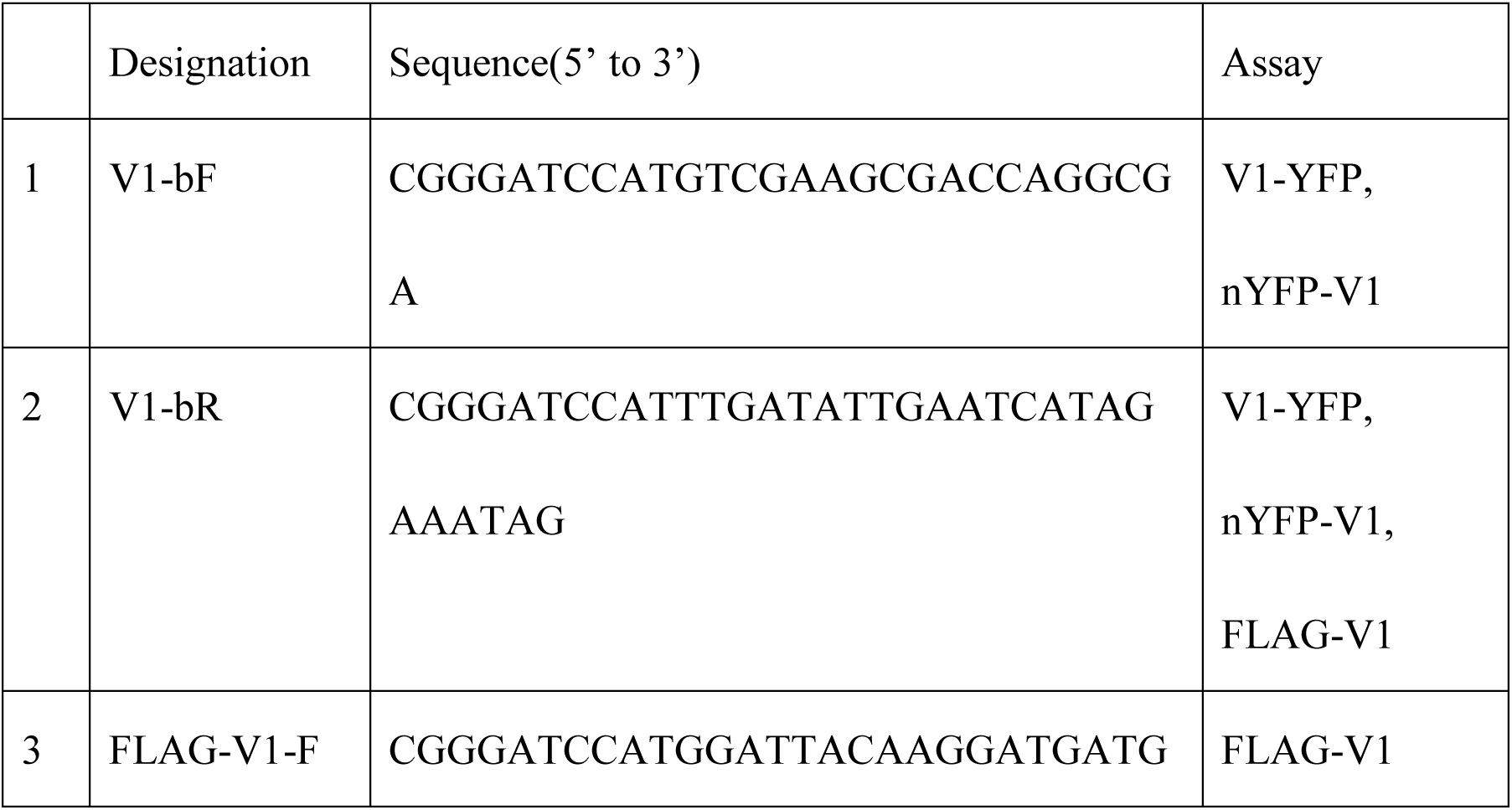
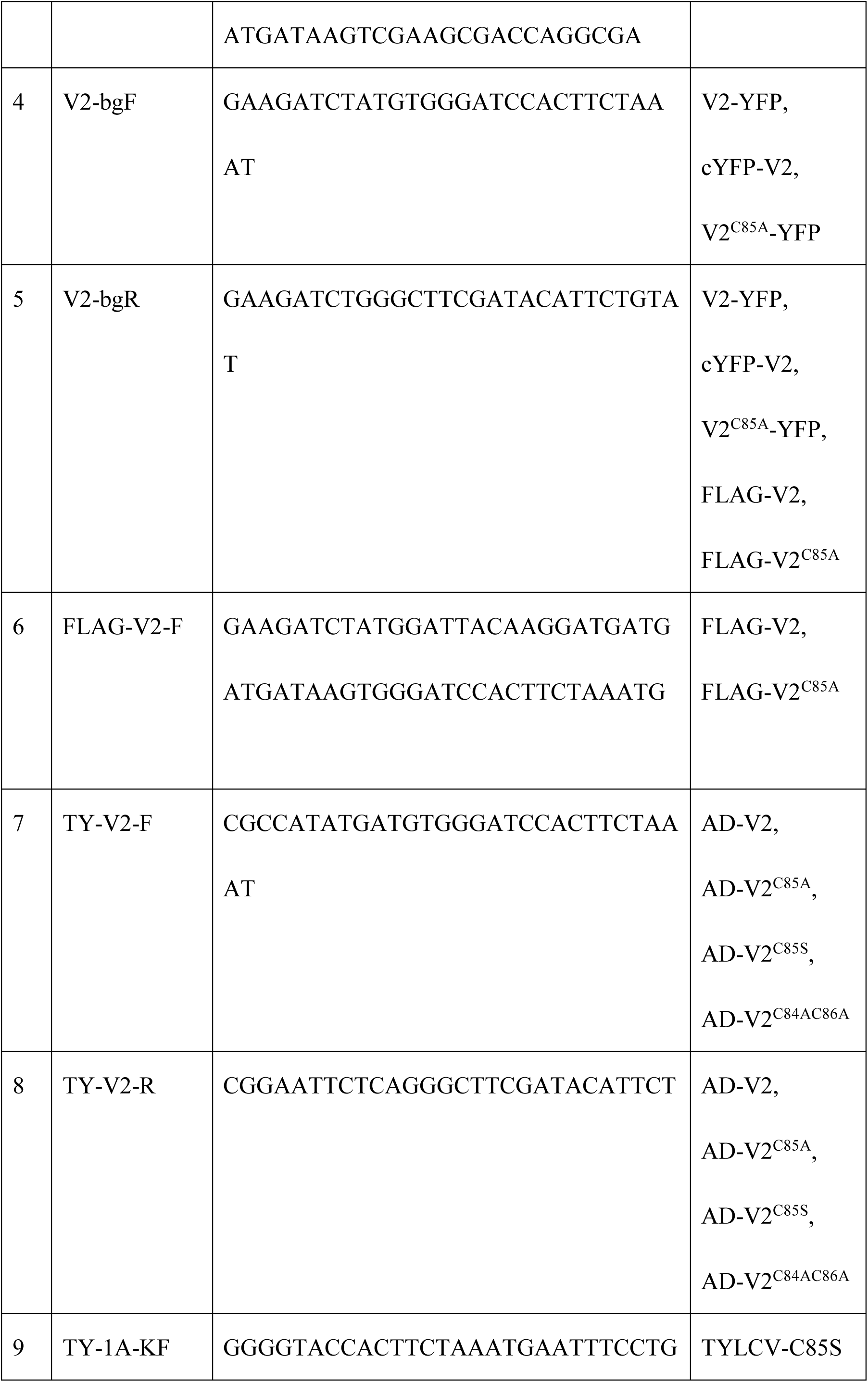
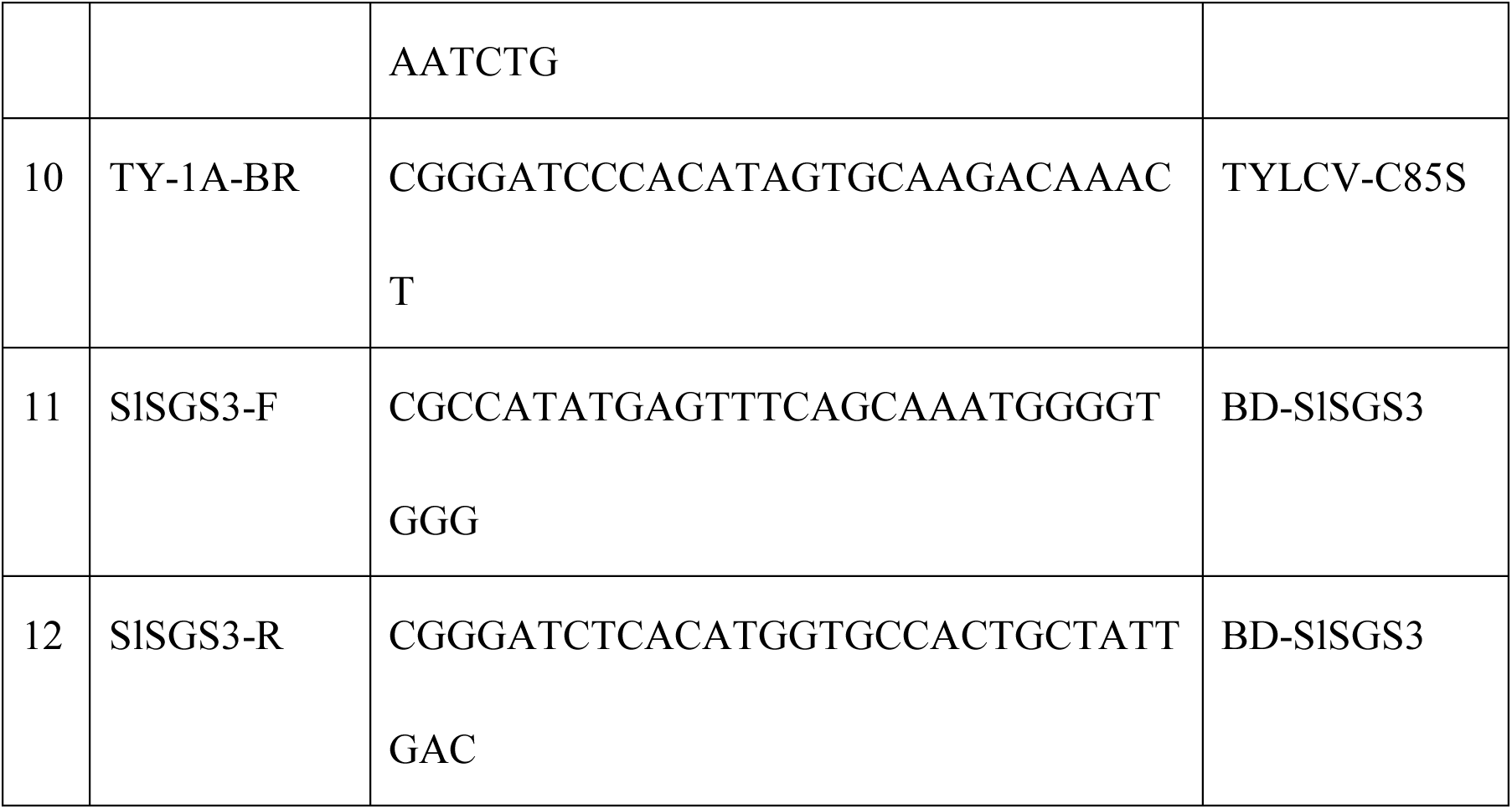
Primers used in this study.

**S1 Fig.**
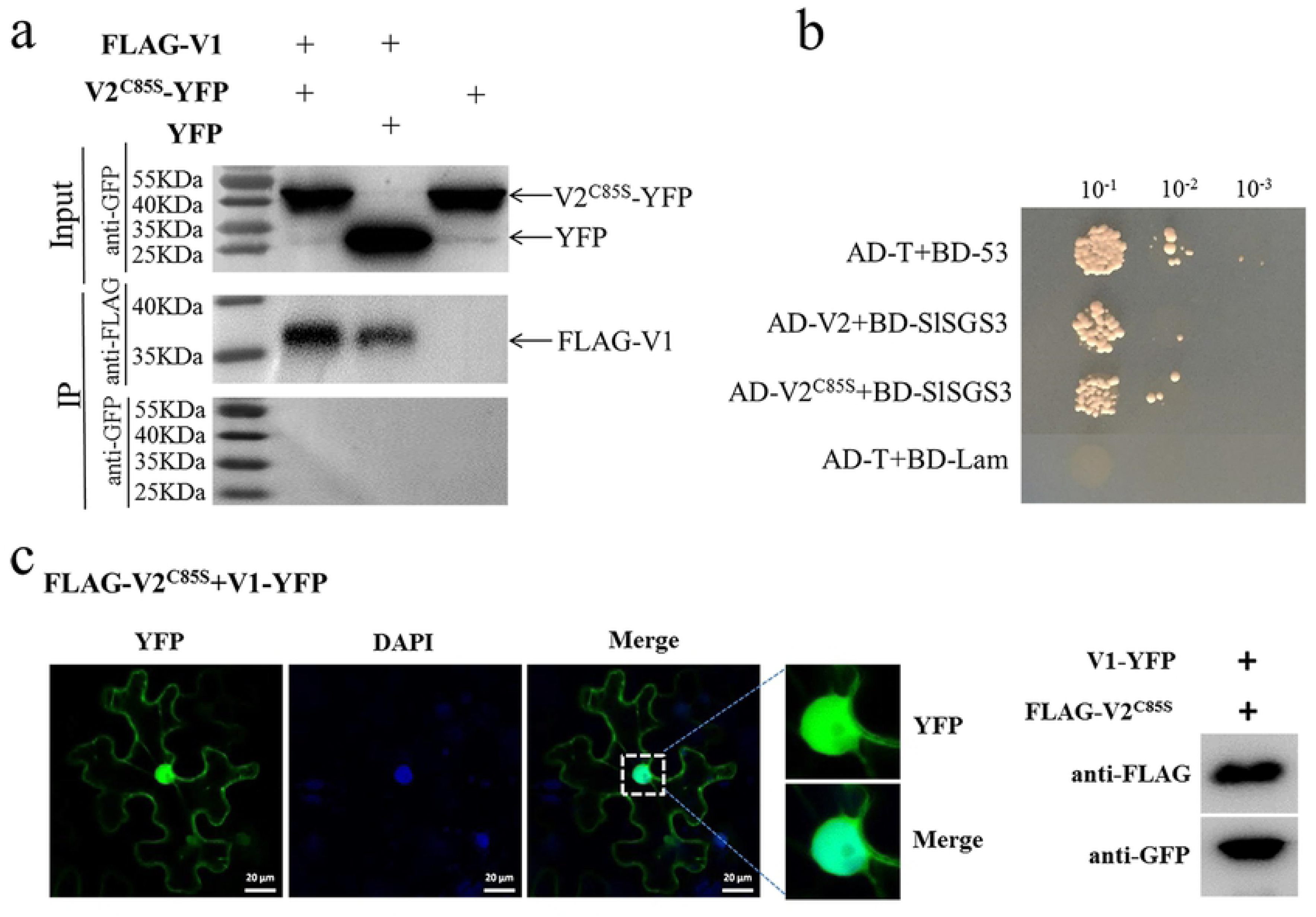
(a) Co-IP assay showing that V2^C85S^ does not interact with V1 protein. The Co-IP assay was performed as in Fig 2a. (b) Analysis of the interaction between SlSGS3 and wt V2 or the V2^C85S^ mutant in the yeast two-hybrid assay. The Y2H assay was performed as in Fig 4c. (c) Subcellular localization of V1 protein that was co-expressed with FLAG-V2^C85S^ in *N. benthamiana* cells. DAPI stains DNA in the nucleus. Bars: 20 µm. Both V1-YFP and FLAG-V2^C85S^ are expressed well as shown by Western blot analysis.

